# Detection and quantification of house mouse *Eimeria* at the species level – challenges and solutions for the assessment of Coccidia in wildlife

**DOI:** 10.1101/636662

**Authors:** Víctor Hugo Jarquín-Díaz, Alice Balard, Jenny Jost, Julia Kraft, Mert Naci Dikmen, Jana Kvičerová, Emanuel Heitlinger

**Author notes:** Note: Supplementary data associated with this article.

## Abstract

Detection and quantification of coccidia in studies of wildlife can be challenging. Therefore, the prevalence of coccidia is often not assessed at the parasite species level in non-livestock animals. Parasite species-specific prevalences are especially important when studying evolutionary questions in wild populations. We tested whether increased host population density increases the prevalence of individual *Eimeria* species at the farm level, as predicted by epidemiological theory. We studied free-living commensal populations of the house mouse (*Mus musculus*) in Germany and established a strategy to detect and quantify Eimeria infections. We show that a novel diagnostic primer targeting the apicoplast genome (Ap5) and coprological assessment after flotation provide complementary detection results increasing sensitivity. Genotyping PCRs confirm detection in a subset of samples and cross-validation of different PCR markers does not indicate a bias towards a particular parasite species in genotyping. We were able to detect double infections and to determine the preferred niche of each parasite species along the distal-proximal axis of the intestine. Parasite genotyping from tissue samples provides an additional indication for the absence of species bias in genotyping amplifications. Three *Eimeria* species were found infecting house mice at different prevalences: *Eimeria ferrisi* (16.7%; 95% CI 13.2 – 20.7), *E. falciformis* (4.2%; 95% CI 2.6 – 6.8) and *E. vermiformis* (1.9%; 95% CI 0.9 – 3.8). We also find that mice in dense populations are more likely to be infected with *E. falciformis* and *E. ferrisi*.

We provide methods for the assessment of prevalences of coccidia at the species level in rodent systems. We show and discuss how such data can help to test hypotheses in ecology, evolution and epidemiology on a species level.

## 1. Introduction

House mice *(Mus musculus)* are the most commonly used mammalian model for biomedical research worldwide (Houdebine, 2004; Vandenbergh, 2013). Laboratory mouse strains are derived mainly from the subspecies *M. m. domesticus* with genetic contributions from other subspecies *(M.m musculus* and *M. m. castaneus)* (Nishioka, 2011; Yang et al., 2007). Establishment of suitable mouse models to better understand infections with coccidia is an ongoing process. Wild rodents and especially wild house mice are an attractive system for first steps in this direction (Ehret et al., 2017).

*Eimeria* (Schneider, 1875) is, with around 1700 species, the most speciose genus in the phylum Apicomplexa (Duszynski, 2011; Perkins et al., 2000). For economical reasons, the most studied parasites in this group are those infecting livestock (Shirley et al., 2005; Su et al., 2003). At least one third of the described species, however, infects rodents (Levine and Ivens, 1965; Zhao and Duszynski, 2001b).

The most commonly used method for detection and identification of coccidia is the flotation and microscopical observation of oocysts shed in faeces during the patent period of infection (Ryley et al., 1976). Unsporulated oocysts, however, are difficult or impossible to differentiate into species (Levine and Ivens, 1988, 1965). Thus, prior to identification the oocyst should be sporulated under specific conditions. In addition, expertise and experience is required for species identification, especially in cases (like ours) of very similar oocyst morphology in different species (Duszynski and Wilber, 1997; Long and Joyner, 1984). For that reason, tools based on DNA amplification and sequencing have been included as routine strategy not only for detection, but also for taxonomic assessment (Fernandez et al., 2003; Hnida and Duszynski, 1999; Kawahara et al., 2010; Morris et al., 2007; Schnitzler et al., 1999; Su et al., 2003; Vrba et al., 2010).

Up to 16 species of *Eimeria* have been described from house mice (Levine and Ivens, 1965) and some of them use different niches in the intestine. The reasons for this diversity are still elusive (Zhao and Duszynski, 2001a) and artificial splitting of morphologically plastic forms of the same species (in the same of different hosts) might contribute to this.

*Eimeria* species described from house mice include *E. falciformis*, the first coccidia described in house mice (Eimer, 1870), which has sometimes been regarded as the most prevalent species in mice (Becker, 1934; Owen, 1976). This species (and especially the BayerHaberkorn1970 isolate) are the most commonly studied coccidia model in laboratory mice. Life cycle progression (Haberkorn, 1970) and host response (Mesfin et al., 1978; Schelzke et al., 2009; Schmid et al., 2012) are relatively well studied and the whole genome of this species has been sequenced and annotated in detail (Heitlinger et al., 2014).

*E. vermiformis* was first described in 1971 (Ernst et al., 1971) but since then, to our knowledge, not reported in wild house mice. Similar to *E. falciformis*, most of the information on this species comes from laboratory infection experiments (Figueiredo-Campos et al., 2018; Rose et al., 1990; Rose and Millard, 1985; Todd Jr and Lepp, 1971), making the timing of life cycle progression and its effect on the host relatively well studied.

*E. ferrisi* was originally described from *M. m. domesticus* from North America (Ankrom et al., 1975; Levine and Ivens, 1965). Laboratory infections with this parasite have confirmed its shorter life cycle, compared to *E. vermiformis* or *E. falciformis* (Schito et al., 1996).

To the best of our knowledge, just few investigation of prevalences and intensities of coccidia has been conducted in free-living populations of *M. musculus* (Ball and Lewis, 1984; Ernst et al., 1971; Golemanski, 1979; Owen, 1976; Parker et al., 2009; Yakimoff and Gousseff, 1938). In the present work we studied the prevalence of *Eimeria* in house mice from a transect of the well-studied European house mice hybrid zone (HMHZ) (Boursot et al., 2003; Ďureje et al., 2012; Phifer-Rixey and Nachman, 2015). We established methods for detection, species identification and quantification of *Eimeria* in these wild commensal populations of house mice.

## 2. Material and methods

### 2.1. Collection of samples

Between 2015 and 2017, 378 house mice *(Mus musculus)* were captured in 96 farms and private properties in a transect 152.27 km long and 114.48 km wide, within the German federal state of Brandenburg (capture permit No. 2347/35/2014) (Figure 1A, Supplementary data 1). On average 20 traps were set overnight per locality. Mice were house individually in cages over night and euthanised by cervical dislocation. All mice were dissected within 24 hours after capture. Faeces for microscopical diagnosis of *Eimeria* spp. were preserved in potassium dichromate (K_2_Cr_2_O_7_) 2.5% (w/v) and stored at 4°C until further processing, colon content was preserved in liquid nitrogen and stored at −80°C. For a subset of 163 mice (from Brandenburg in 2016) tissue samples from cecum and ileum were collected for DNA extraction and molecular identification of *Eimeria* spp. All samples were kept in liquid nitrogen during transportation and maintained at −80°C until processing.

**Figure 1.**
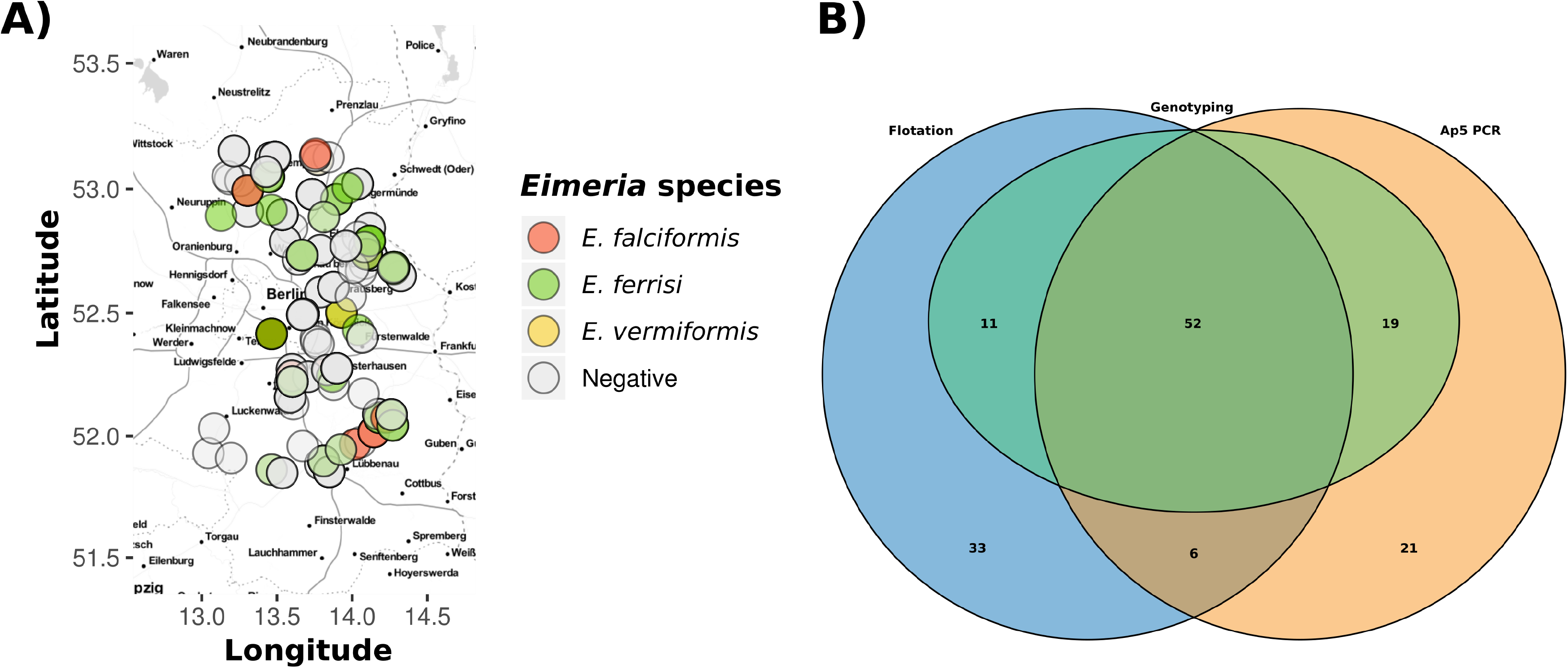
Geographical localization of house mice *(Mus musculus)* collected for this study and comparison of diagnostic methods for *Eimeria*. A) Localization from the 378 mice included in the present study, colors indicate the *Eimeria* species identified for each. B) Venn diagram showing the overlap between detection methods and successful genotyping identification of the isolates.

### 2.2. Flotation and microscopical analysis of oocyst

Fecal samples were washed with tap water to eliminate potassium dichromate and homogenized. After oocyst were flotated using a saturated salt solution (specific gravity = 1.18—1.20), they were collected by centrifugation (3,234 × g/room temperature/ 10 minutes), washed with distilled water (1,800 × g/room temperature/10 minutes). The flotations were screened for the presence of oocyst using a Leica^®^ DM750 M light microscope under the 10X objective. To estimate the intensity of infection, flotated oocysts were counted using a Neubauer chamber and the results were expressed as oocyst per gram (OPG) of faeces. Samples were then preserved in a fresh solution of potassium dichromate 2.5% (w/v) and sporulated in a water bath at 30°C for 10-12 days for further characterisation.

*Eimeria* isolates, corresponding to different phylogenetic groups (see below), were selected to take photomicrographs of sporulated oocysts using a Carl-Zeiss microscope at 100x magnification. Measurements were made on ~30 oocysts and ~30 sporocysts, using Adobe Photoshop CC v14.2.1 (3778 pixels = 100, 0000 μm). Length and width were measured and reported in micrometers. The Length/Width (L/W) ratio was calculated for both oocysts and sporocysts including means, standard deviation and variation coefficients. Additionally, main morphological traits (oocyst wall, oocyst residuum, micropyle, polar granule, sporocyst residuum, refractile bodies and Stieda body) were described, according to the protocol of Duszynski and Wilber (1997).

### 2.3. DNA extraction

DNA from colon content was extracted using the NucleoSpin^®^ Soil kit (MACHEREY-NAGEL GmbH & Co. KG, Düren, Germany) according to the manufacturer’s recommendations, adding a mechanical lysis process in a Mill Benchtop Mixer MM 2000 (Retsch GmbH, Haan, Germany). DNA from cecum and ileum tissues was isolated using the innuPREP DNA Mini Kit (Analytik Jena AG, Jena, Germany) following the instructions of the manufacturer after disruption of the tissue with liquid nitrogen in a mortar. Quality and quantity of isolated DNA was measured spectrophotometrically in a NanoDrop 2000c (Thermo Scientific, Waltham, USA).

### 2.4. PCR amplification for detection (ap tRNA) and identification (nu 18S rDNA and mt COI)

For detection of *Eimeria*, amplification of a conserved tRNA region of the apicoplast genome (Ap5) was used. Primers Ap5_Fwd (YAAAGGAATTTGAATCCTCGTTT) and Ap5_Rev (YAGAATTGATGCCTGAGYGGTC) were designed based on the complete apicoplast genomes sequences available in the GenBank from *Eimeria tenella* (NC_004823.1), *E. falciformis* (CM008276.1) and *E. nieschulzi* (JRZD00000000.1).

For all samples with oocysts detected during flotation or successful amplification of Ap5, genotyping PCRs were performed to confirmation of detection and further identification of parasite species. A fragment of nuclear small subunit ribosomal RNA (~1,500 bp) and mitochondrial cytochrome C oxidase subunit I (~ 800 bp) were amplified using primers 18S_EF and 18S_ER (Kvičerová et al., 2008) and Cocci_COI_For/Rev (Ogedengbe et al., 2011), respectively. An alternative pair of primers was used in case of failure to amplify COI: Eim_COI_M_F (ATGTCACTNTCTCCAACCTCAGT) and Eim_COI_M_R (GAGCAACATCAANAGCAGTGT). These primers were designed based on the mitochondrial genome of *E. falciformis* (CM008276.1) (Heitlinger et al., 2014) and amplify a ~700 bp fragment of COI gene.

PCR reactions were carried out in a Labcycler (SensoQuest GmbH, Göttingen, Germany) using 0.025 U/μL of DreamTaqTM DNA Polymerase (Thermo Scientific, Waltham, USA), 1X DreamTaq Buffer, 0.5 mM dNTP Mix, 0.25 μM from each primer and 1—20 ng/μL of DNA template in 25 μL reaction. A concentration of 0.25 mM dNTP Mix and a supplementation with 2 mM MgCl_2_ was used for the amplification of Ap5. The thermocycling protocol consist of 95 °C initial denaturation (4 min) followed by 35 cycles of 92 °C denaturation (45 s), annealing at 52°C (30 s/Eim_COI); 53 °C (45 s/18S_E); 55 °C (30 s/Cocci_COI); 56 °C (30 s/Ap5); 72 °C extension 90 s (18S_E), 20 s (Cocci_COI/Eim_COI) or 45s (Ap5) and a final extension at 72 °C (10 min). DNA from oocyst of *E. falciformis* BayerHaberkorn1970 strain and DNA from colon content of a non-infected NMRI mouse were used as positive and negative controls, respectively.

All PCR products with the expected size were purified using the SAP-Exo Kit (Jena Bioscience GmbH, Jena, Germany) and Sanger sequenced from both directions by LGC Genomics (Berlin, Germany). Quality assessment and assembly of forward and reverse sequence was performed in Geneious v6.1.8. All sequences were submitted to NCBI GenBank (Accession numbers: nu 18S rDNA [MH751925—MH752036] and mt COI [MH777467—MH777593 and MH755302— MH755324] (Supplementary data S2).

### 2.5. Eimeria detection in tissue by qPCR

For mice collected in 2016 *(n=* 163) cecum and ileum tissue was screened using qPCR. Primers targeting a short fragment of mt COI were used to amplify DNA from from intracellular stages of *Eimeria* (Eim_COI_qX-F, TGTCTATTCACTTGGGCTATTGT; Eim_COI_qX-R GGATCACCGTTAAATGAGGCA). Amplification reactions with a final volume of 20 μL contained 1X iTaqTM Universal SYBR^®^ Green Supermix (Bio-Rad Laboratories GmbH, München, Germany), 400 nM of each primer and 50 ng of DNA template. Cycling in a Mastercycler^®^ RealPlex 2 (Eppendorf, Hamburg, Germany) was performed with the following program: 95 °C initial denaturation for 2 min, followed by 40 cycles of denaturation at 95 °C for 15 s, annealing at 55 °C for 15 s and extension 68 °C for 20 s. Melting curves were analysed to detect eventual primer dimer formation or non-specific amplification. As internal reference for relative quantification the CDC42 gene from the nuclear genome of the house mouse was amplified (Ms_gDNA_CDC42_F CTCTCCTCCCCTCTGTCTTG; Ms_gDNA_CDC42_R TCCTTTTGGGTTGAGTTTCC). Infection intensity was estimated as the ΔCt between mouse and *Eimeria* amplification (CtMouse-*CtEimeria*). To correct for background noise a detection threshold was estimated at ΔCt = −5 and only results above this value were considered infected. ΔCtIleum and ΔCtCecum were compared for samples above the threshold in both tissues to assess primary tissue occurrence (Ahmed et al., 2019). In samples positive for qPCR, *Eimeria* genotyping was performed based on DNA extracted from tissue, as described above (see *2.4*).

### 2.6. Molecular identification of Eimeria spp. isolates: 18S and COI phylogenetic analysis

As strategy for molecular identification, datasets of nu 18S and mt COI sequences were compiled. Sequences generated for the present work were compared to databases sequences using NCBI BLAST and most similar sequences were selected. Based on this, sequences for *E. falciformis, E. vermiformis* and *E. ferrisi* were downloaded from GenBank as a reference. COI sequences were aligned by translation using the Multiple Align algorithm and translation frame 1 with the genetic code for “mold protozoan mitochondrial”, 18S sequences were aligned using MUSCLE (Edgar, 2004), both through Geneious v6.1.8.

Phylogenetic trees for all datasets were constructed using Maximum Likelihood (ML) and Bayesian inference (BI) methods, implemented in PhyML v3.0 (Guindon et al., 2010) and MrBayes v3.2.6 (Huelsenbeck and Ronquist, 2001; Ronquist et al., 2012), respectively. The most appropriate evolutive models for each datasets were determined in JModelTest v2.1.10 (Posada, 2008). For ML trees, a bootstrap analysis with 1000 replicates was performed, whereas MCMC for BI was run with two cold and two hot chains for 1,000,000 generations or until the average split frequency was below 0.05. The concatenated dataset was analysed using partitions and locus-specific models. Visualization of the trees was done with FigTree v1.4.2 (Rambaut, 2012).

### 2.7. Statistical analysis

All statistical analyses were performed in R (R Development Core Team, 2008). Prevalence of *Eimeria* was calculated as the proportion of positive samples in the total number of analysed samples. The 95% confidence interval [CI 95%] was calculated using the Sterne’s exact method (Sterne, 1954) implemented in the package “epiR” v0.9-99 (Nunes et al., 2018). Prevalences were tested for statistical differences with the Fisher’s exact test (Fisher, 1922).

To assess the significance in primer bias, logistic regression models were used to estimate the probability to successfully amplify and sequence a specific genetic marker for each *Eimeria* species. The response variable in these models was the amplification and sequencing success with a particular primer pair (COCCI_COI_F/R, Eim_COI_M_F/R or 18S_E F/R), and the predictors were the species identity (as determined with the other markers only, to make response and predictors independent) and additionally the detection of an infection with Ap5 and Flotation. These models were fitted first for COCCI_COI_F/R as response, then for the combined probability of successful COI genotyping and finally for 18S genotyping as response. Tables were produced for the summary of models using the package “jtools” v2.0.0 (Long, 2019).

For each *Eimeria* species logistic regression models were used to test whether the infection is influenced by host density or by the presence of the other two *Eimeria*. We use as response variable the infection status by *E. ferrisi, E. falciformis* or *E. vermiformis*, independently, and the total number of mice cough per locality per year and the infection status by a different *Eimeria* species as predictors.

Differences on oocyst and sporocyst L/W ratios between *Eimeria* species were tested for significance with an analysis of variance fitting a linear model using the species assignment as predictor with a Tukey HSD post hoc test adjusting for multiple comparisons.

## 3. Results

### 3.1. *Sampling and Eimeria* spp. *detection*

We used flotation of oocyst from faeces and PCR amplification of a novel diagnostic maker (Ap5) from colon content DNA to detect *Eimeria* parasites in a total of 378 house. Overall prevalence was 25.9% [95% CI = 21.7 – 30.7] (98/378) for PCR and 27.0% [95% CI = 22.7 – 31.7%] (102/378) for flotation. These estimates are not significantly different (Fisher exact test, p> 0.05). However, both techniques considered together estimate a higher prevalence of 37.6% [95% CI = 32.8 – 42.6] (142/378), meaning that 44 and 40 positive results were detected only by flotation or PCR, respectively (Fig. 1B). We further aimed to provide species specific identification and to consolidate results from the two different detection methods.

### 3.2. Molecular identification of Eimeria isolates – (phylogenetic analysis nu 18S and mt COI)

*Eimeria* species were identified by phylogenetic analysis of nu 18S and mt COI sequences, the most commonly used molecular markers of apicomplexan parasites. To identify our isolates, sequences were compared with references from the NCBI database. Sequences from three previously described *Eimeria* species infecting *M. musculus* showed highest BLAST similarities and phylogenetic clustering. This approach ignores the problem whether isolates from different hosts would be assigned to the same phylogenetic clusters while they are regarded different species by taxonomists.

The nu 18S phylogenetic tree was inferred based on 80 sequences (540—1,795 bp), 73 of them from wild house mice generated in our study (3 from ileum tissue, 16 from cecum tissue and 54 from colon content, see below). *Eimeria* species previously described in house mouse were represented by *E. falciformis* (AF080614), *E. vermiformis* (KT184355) and *E. ferrisi* (KT360995). In addition, one newly generated sequence from *E. falciformis* strain BayerHaberkorn1970 (MH751998) was also included. Sequence identity of our isolates to this references sequences was above 98% and even 100% in most of the cases for this marker. *Isospora* sp. sequences identified in *Talpa europaea* moles were used as outgroup. Both ML and BI rooted trees shared a topology placing our sequences at the same positions in relation to reference sequences with high support (bootstrap values and posterior probabilities are shown in Fig. 2A). The sequences clustered in three well supported monophyletic groups (Fig. 2A).

**Figure 2.**
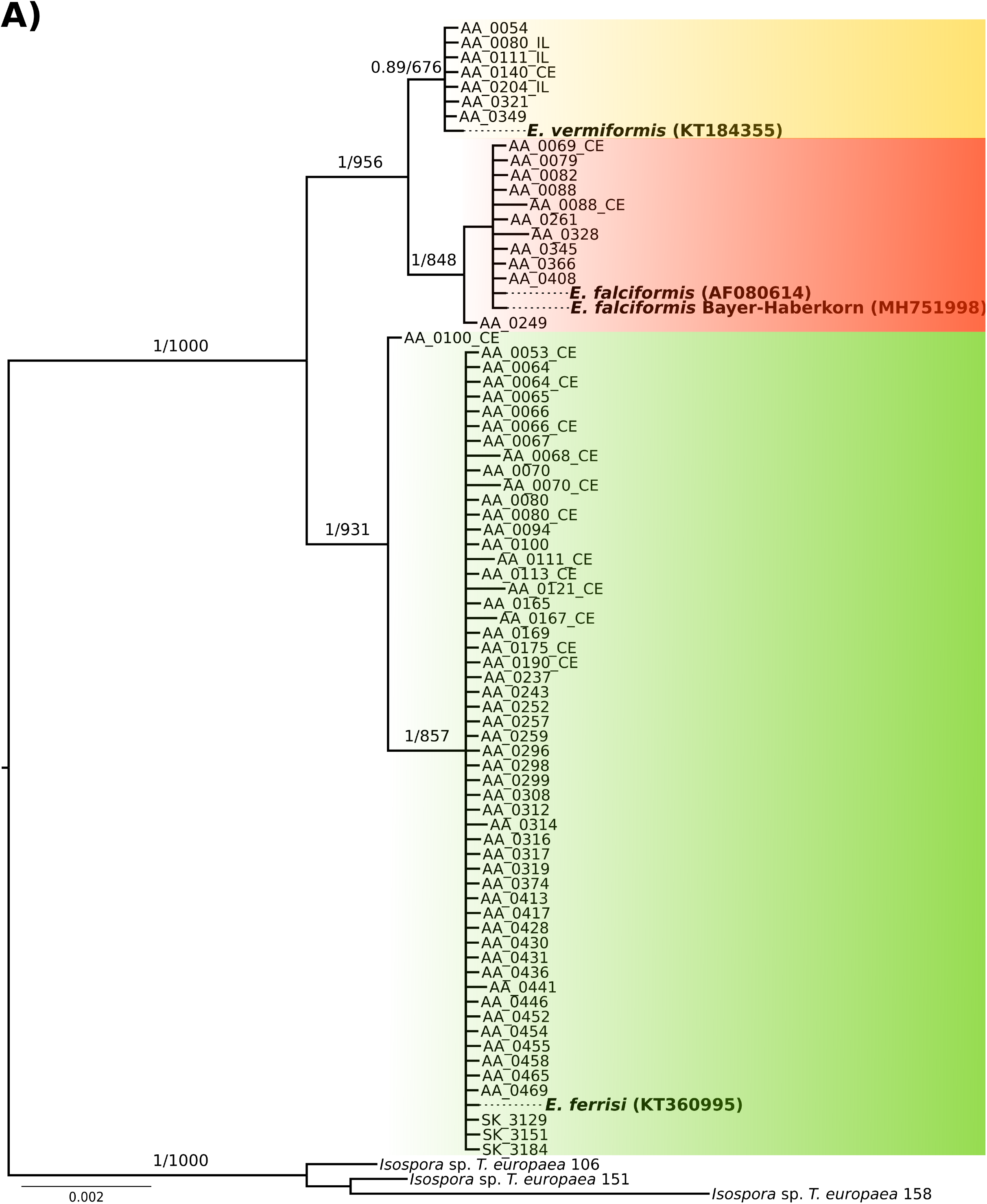
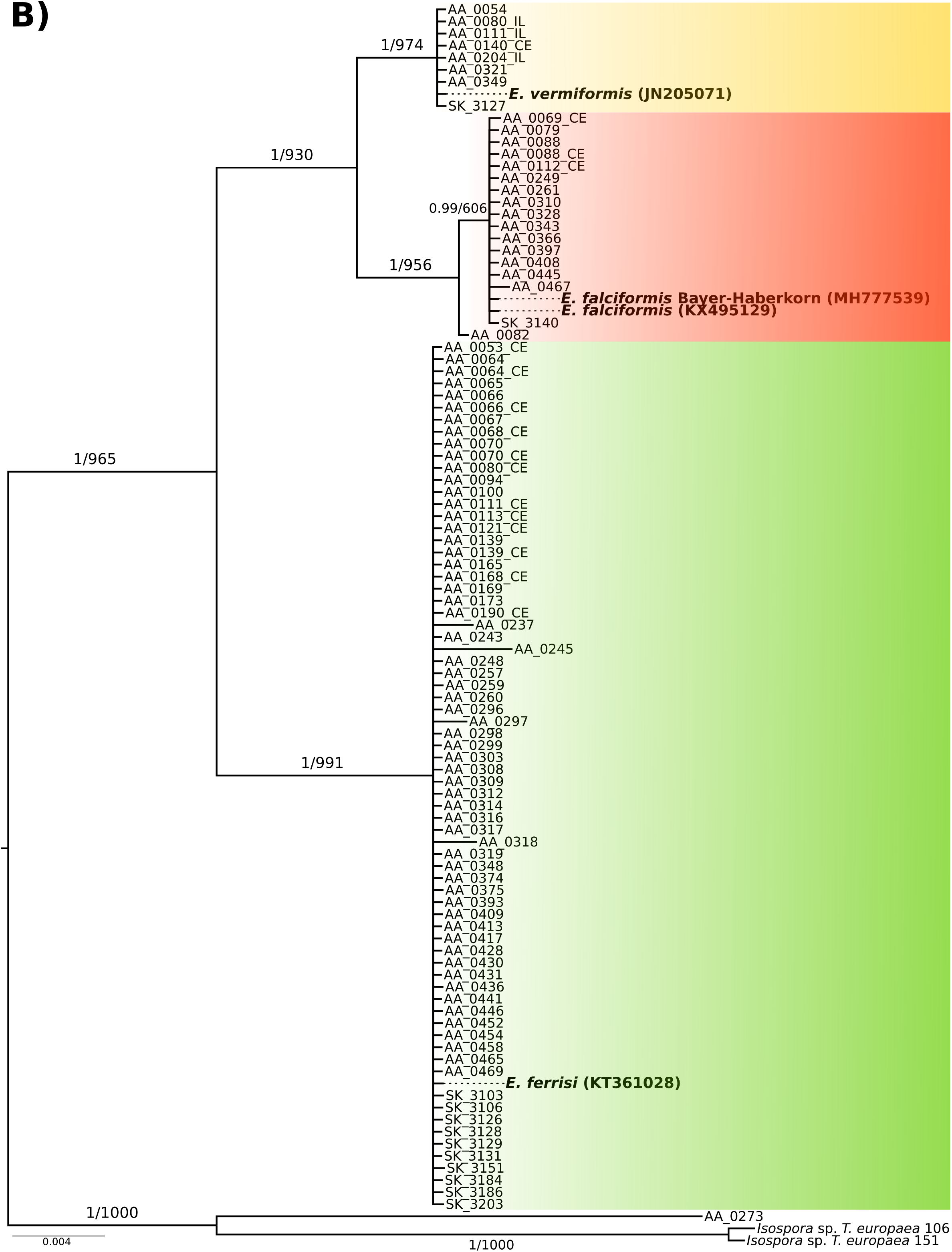
Phylogenetic trees based on 18S rRNA and COI sequences. Sequences of 18S A) and COI B) were used to infer the molecular identification of wild-derived isolates of *Eimeria*. In both phylogenies, our isolates clustered in three groups one close to *E. falciformis* (red), other close to *E. ferrisi* (green) and finally one to *E. vermiformis* (yellow). Numbers in the branches represent the Bayesian posterior probability and the non-parametric bootstrap value. In bold are indicate the reference sequences for each species. CE and IL make reference to sequence derived from cecum or ileum tissue DNA, respectively.

The phylogenetic tree for mt COI was based on 103 sequences (519—804 bp), 97 of which were obtained from *Eimeria* infecting wild house mice (3 from ileum, 16 from cecum tissue and 78 from colon content). Reference sequences from house mouse *Eimeria (E. ferrisi*, KT361028; *E. falciformis*, KX495129 and MH777539; *E. vermiformis* JN205071) identified by BLAST searches showed an identity of above 98% to respective groups of our isolates. We defined *Isospora* sp. from *Talpa europaea* as outgroup for rooting. ML and BI rooted trees based on alignments of these COI sequences shared a general topology with respect to the placement of our isolates in relation to reference sequences. Bootstrap values and posterior probabilities for support of these placements are shown in Fig. 2B. The sequences derived from house mice cluster in three monophyletic groups including reference sequences for *E. falciformis (n=* 17, sequences from our study), *E. ferrisi (n=* 72) and *E. vermiformis (n=* 8) (Fig 2B). Phylogenies based on concatenated supermatrices for the two markers show the same topology concerning placement of isolates from the present study (Supplementary data S3 and S4).

### 3.3. Morphometrical and morphological comparison of oocysts

For further support assignment of the three phylogenetic groups of *Eimeria* from house mouse, we characterized sporulated oocysts morphologically (Table 1). *E. falciformis, E. ferrisi* and *E. vermiformis* oocyst shared most of the traits we evaluated and showed overlapping morphometry (Fig. 3A). The length/width ratio of *E. vermiformis* oocysts, however, was significantly higher (1.29; 95%CI = 1.26—1.33; n= 35) than that of *E. falciformis* (1.17; 95%CI = 1.14—1.20; n= 31) and *E. ferrisi* oocysts (1.23; 95%CI = 1.21—1.25; n= 127) (Tukey HSD, *p* <0.05) (Fig. 3B; Supplementary data S5). This means that *E. vermiformis* has more ellipsoidal oocysts than the other two species. Other morphological characteristics of oocysts (smooth wall, absence of micropyle, presence of polar granule and absence of oocyst residuum) are very similar or identical between the three species (Table 1).

**Figure 3.**
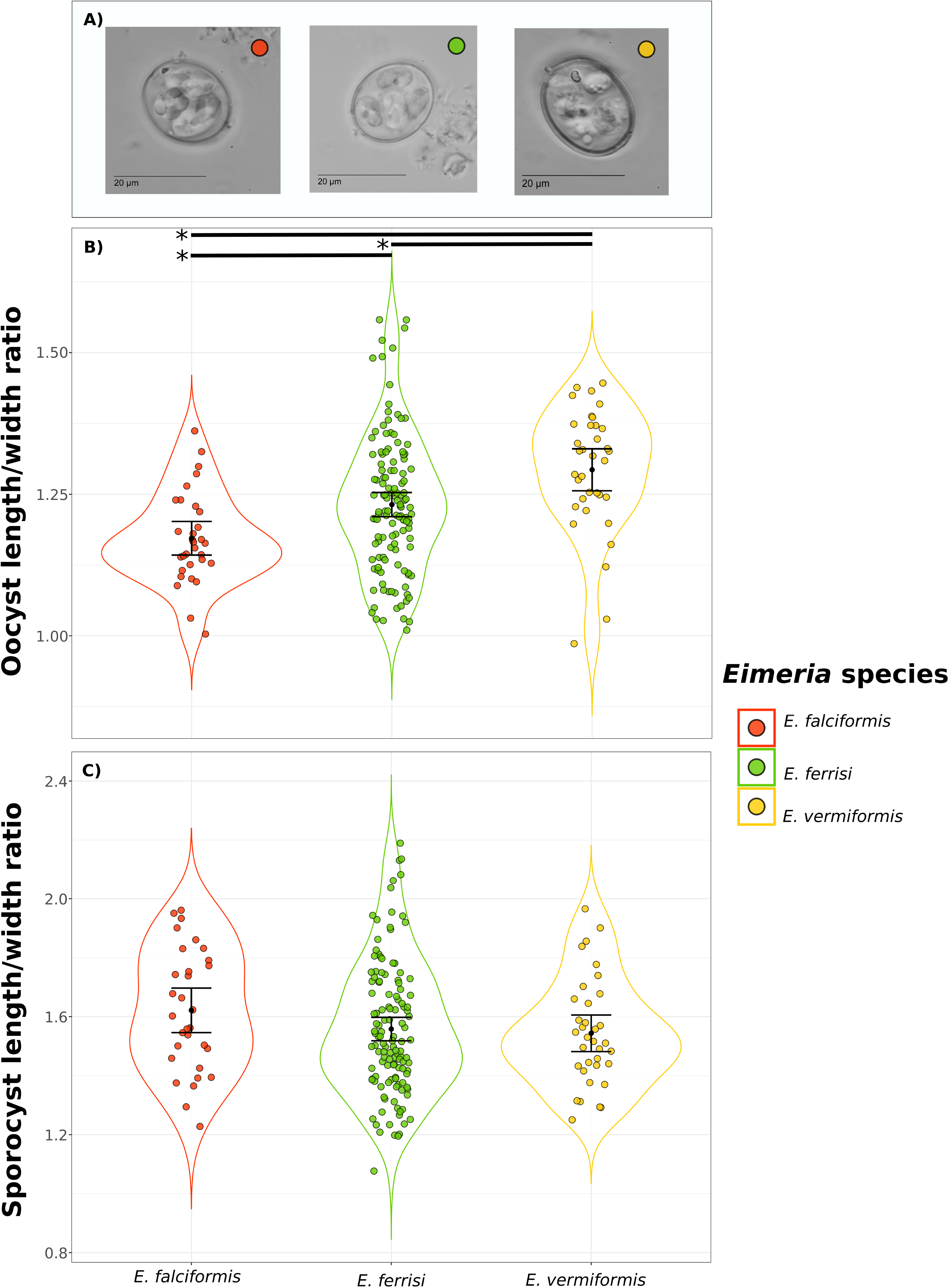
Morphological and morphometrical characteristics of *Eimeria* oocyst isolated from *Mus musculus*. a) Photomicrographs at 1000x amplification of *Eimeria* oocyst from the three species isolated from *Mus musculus* (red = *E. falciformis;* green = *E. ferrisi* and yellow = *E. vermiformis*). Length/Width ratio from b) oocyst and c) sporocysts corresponding to each species *(E. falciformis n=* 31; *E. ferrisi n=* 127 and *E. vermiformis* n= 35). Mean +/− 95% Confidence Interval is plotted. * Represent significant difference (Tukey HSD, p<0.05).

**Table 1.**
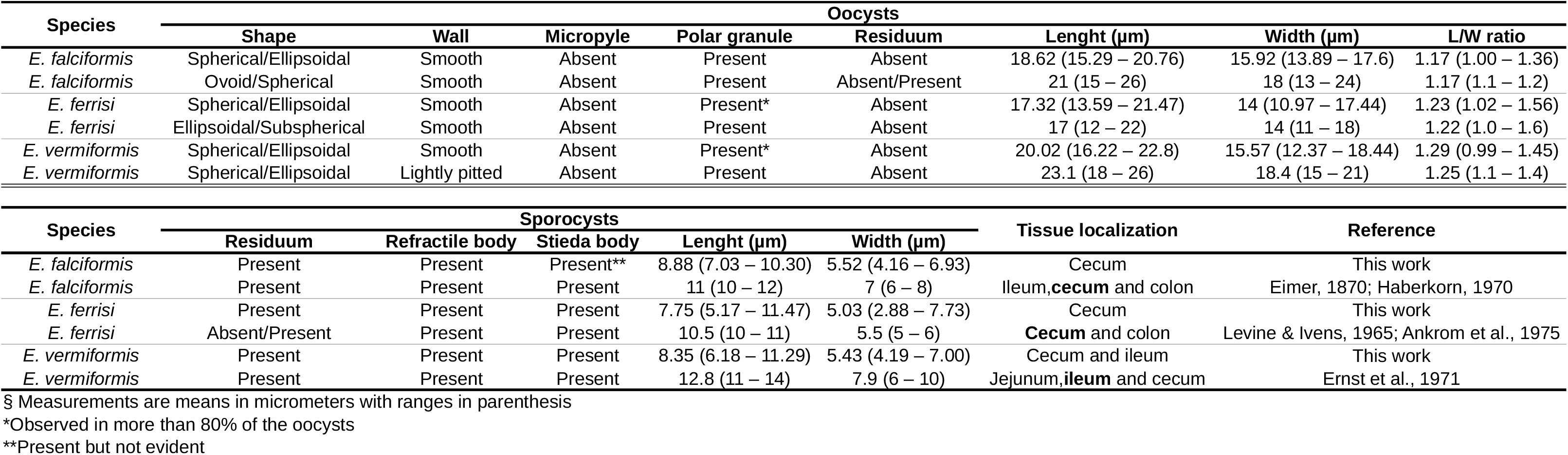
Morphological and morphometrical characteristics from *Eimeria* wild-derived isolates and reference morphotypes^3^.

Morphological measurements of sporocysts are not significantly different between the three species found in house mice (Fig. 3C). We also observed the presence of a sporocyst residuum, refractile bodies and Stieda bodies in all species uniformly, in agreement with previous descriptions (Ankrom et al., 1975; Eimer, 1870; Ernst et al., 1971; Haberkorn, 1970).

### 3.4. Proximal-distal occurrence of infection and double infections

We detected DNA from endogenous stages by qPCR in 27 of 163 samples analysed (Supplementary data S6). We differentiate detection between small and large intestine, analysing ileum as the most distal tissue of the small intestine and cecum as the most proximal tissue of the large intestine. Detection was either limited just to cecum (*n* = 19), to ileum (*n* = 2) or possible in both tissues (*n* = 6). Infections in cecum were identified as *E. falciformis* (*n* = 4), *E. ferrisi* (*n* = 17) and *E. vermiformis* (*n* = 1) Detections in ileum (*n* = 2) were identified as *E. vermiformis*. In two mice positive in both tissues, it was possible to identify *E. ferrisi* in cecum and *E. vermiformis* in ileum, providing evidence that these animals presented a double infection (that is, simultaneous infections with different isolates; Fig. 4).

**Figure 4.**
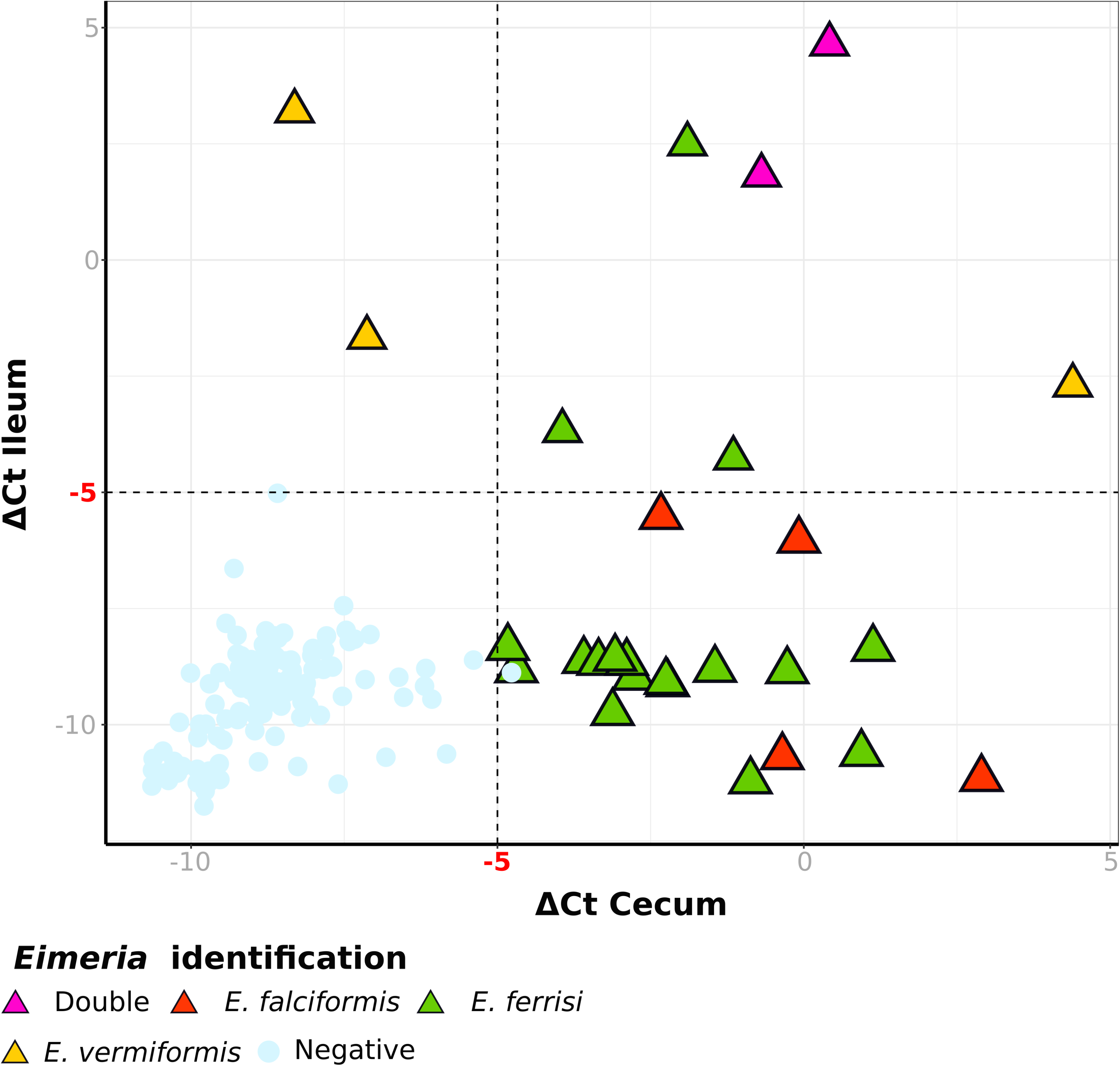
qPCR detection of intracellular stages of *Eimeria* in cecum and ileum from *Mus musculus*. -Delta Ct value (Ct_Eimeria_ – Ct_Mouse_) from each tissue for 164 mice are plotted on the graph. The dotted line indicate the threshold of −5, values above the line are considered positive for the corresponding tissue. Circles represent negative samples, triangles indicate samples with *Eimeria* species identification and colors correspond to the *Eimeria* species identified in those samples.

### 3.5. Amplification efficiency of different markers

As *Eimeria* detection by PCR and the determination of species identity could be biased especially in cases of double infections, we analysed differences in amplification efficiency of the three primer pairs used for the molecular identification of *Eimeria* species (Table 2). In a cross-validation approach we compare the likelihood to amplify a maker given the species identification with the other marker. The amplification and sequencing efficiency of Cocci_COI primer pair was significantly higher for *E. ferrisi* isolates (logistic regression, *p* < 0.001) than for *E. falciformis* isolates (the letter determined by the 18S marker). Using the novel primer pair Eim_COI_M_F/R the sequencing results were complemented and we detected no significant differences in the combined amplification efficiency for both primers (logistic regression; *p* = 0.62).

**Table 2.** Statistical models used to analyse primer preference to different *Eimeria* species.

| Predictors | Model 1 (Cocci_COI) | Model 2 (Cocci_COI + Eim_COI) | Model 3 (18S_EF/R) |
| --- | --- | --- | --- |
| Intercept | -5.80 ***<br>(1.11) | -1.55<br>(1.27) | -1.40<br>(0.87) |
| Ap5 | 3.50 ***<br>(0.59) | 3.99 ***<br>(0.60) | 0.78<br>(0.62) |
| Flotation | 1.64 **<br>(0.52) | 2.15 ***<br>(0.54) | 1.59 **<br>(0.51) |
| Identification *E. ferrisi* (other marker) | 5.10 ***<br>(1.14) | 0.28<br>(1.35) | 0.32<br>(0.63) |
| Identification *E. vermiformis* (other marker) | 17.23<br>(1385.38) | 11.98<br>(1385.38) | 1.71<br>(1.33) |
| No identification (other marker) | 0.97<br>(0.92) | -3.57 **<br>(1.23) | -3.44 ***<br>(0.87) |
| Identification *Eimeria* spp. (COI) |  |  | -14.80<br>(1455.40) |
| N | 378 | 378 | 378 |
| AIC | 128.09 | 119.18 | 143.39 |
| BIC | 151.70 | 142.79 | 170.93 |
| Pseudo R2 | 0.77 | 0.82 | 0.70 |

The upper number represents the estimate and numbers in brackets represent standard error for each predictor.
Intercept is *Eimeria falciformis* identification with other marker
Other marker refers to 18S based identification for COI models or vice versa
N (Total number of samples), AIC (Akaike information criterion), BIC (Bayesian information criterion).
\*\*\* p < 0.001; \*\* p < 0.01; \* p < 0.05.

Similarly, we did not detect significant differences in the probability to obtain an 18S sequence (using the 18S_EF/R primer pair) between *Eimeria* species as determined by (combined) COI assessment (logistic regression; *p* = 0.25). Differences in PCR efficiency for Cocci_COI make it likely that markers amplify different species in case of double infections in a single isolate. Both detection by flotation and diagnostic (Ap5) PCR significantly increase the likelihood amplification for all primers (COI or 18S). The statistical models for biased amplification hence also confirm that both detection methods provide complementary results, even while controlling for detection with the other method (Table 2).

### 3.6. Prevalence of the different Eimeria species

Genotyping amplification and sequencing with either 18S or COI primers (or both) was possible for samples in which infections had been detected (n=82) (Fig. 1B). Both COI and 18S genotyping PCRs hence support detection by flotation and diagnostic PCRs in this subset of samples. Furthermore amplification of both or either one maker was fully sufficient to assign the isolate to an *Eimeria* species (Fig. 2), allowing us to resolve prevalence on the species level.

These corrections and controls allow us to determine prevalence at the species level: *E. ferrisi* was identified at a higher prevalence of 16.7% (63/378, 95%CI = 13.2 — 20.7) in comparison to *E. falciformis* (16/378, 4.2% [95%CI = 2.6—6.8]) and *E. vermiformis* (7/378, 1.9% [0.9—3.8]). Considering prevalence at the level of farms, *E. ferrisi* was detected in 29.2% (28/96, 95%CI = 20.7 – 39.0), *E. falciformis* in 12.5% (12/96, 95%CI = 7.1 – 20.7) and *E. vermiformis* in 7.3% (7/96, 95%CI = 3.5 – 14.4) of sampled localities. 25 (of in total 96) farms, had mice with single *Eimeria* species detected, and 10 had more than one species detected. In all cases *E. ferrisi* was detected (5 farms with *E. ferrisi – E. falciformis*, 3 farms with *E. ferrisi – E. vermiformis* combination, and 2 farms with the three species). Mice presenting double infections were caught at farms at which infections with the both *Eimeria* species was found in other mice independently.

We used the number of mice caught per farm as a proxy for population density, assuming roughly equal trapping effort at all localities. We then question whether population density affects prevalence by testing differences in the likelihood of a mouse individual to be infected dependent on that population density. We detect that the likelihood of infection is significantly increased for both *E. ferrisi* and for *E. falciformis* (logistic regression, *p* < 0.05; Table 3). Infection with *E. falciformis* got more likely by 19%, infection with *E. ferrisi* by 14% with each mouse caught at the same locality. We also included the detection of other *Eimeria* species in the model for each species and did not find a significant influence (p > 0.05) on likelihood of infection.

**Table 3.** Statistical models used to analyse factors influencing infection to differentE/mer/aspecies.

| Predictors | Model 1 (*E. ferrisi*infection) | Model 2 (*E. falciformis*infection) | Model 1 (*E. vermiformis*infection) |
| --- | --- | --- | --- |
| Intercept | -1.62 ***<br>(0.35) | -3.32 ***<br>(0.61) | -3.99 ***<br>(0.83) |
| Total caught mice | 0.13 *<br>(0.06) | 0.18 *<br>(0.08) | 0.11<br>(0.10) |
| *E. ferrisi*infection |  | 0.92<br>(0.70) | 0.29<br>(1.06) |
| *E. falciformis*infection | 0.92<br>(0.69) |  | 1.60<br>(0.92) |
| *E. vermiformis*infection | 1.61<br>(0.91) | 0.41<br>(1.03) |  |
| N | 104 | 104 | 104 |
| AIC | 120.99 | 71.61 | 52.03 |
| BIC | 131.57 | 82.18 | 62.61 |
| Pseudo R2 | 0.18 | 0.19 | 0.17 |

The upper number represents the estimate and numbers in brackets represent standard error for each predictor.
N (Total number of samples), AIC (Akaike information criterion), BIC (Bayesian information criterion).
\*\*\* p < 0.001; \*\* p < 0.01; \* p < 0.05.

## 4. Discussion

In this study we identify *Eimeria* species in wild commensal populations of house mice *(Mus musculus*). We show that detection and identification of this group of rodent coccidia can be challenging and propose to complement classical coprological assessment with molecular tools: a highly sensitive detection PCR, genotyping PCRs for species identification and qPCR for localization and detection of double infections. Based on this we identified three different *Eimeria* species in the house mouse: *E. ferrisi, E. falciformis* and *E. vermiformis*. Morphological characteristics and preferential occurrence were congruent with the assignment of isolates to the above species. We use our results to show a positive effect on host density on prevalence of *E. ferrisi* and *E. falciformis*.

Few studies report prevalences of *Eimeria* in wild populations of *Mus musculus*. Prevalences range from 3% to 40% for isolates classified either as *E. falciformis, E. ferrisi* or *E. vermiformis* (Ball and Lewis, 1984; Ernst et al., 1971; Golemanski, 1979; Levine and Ivens, 1965; Tattersall et al., 1994). Other studies make no assessment at the species level (detection as *Eimeria* spp) (Moro et al., 2003; Owen, 1976; Parker et al., 2009; Yakimoff and Gousseff, 1938).

A recent study in rodents (other then *Mus musculus*) in central Europe reported an *Eimeria* spp. prevalence of 32.7% based on coprological observations (Mácová et al., 2018). At 37.6% the overall prevalence for all *Eimeria* species in our study in house could be considered high in comparison to all these studies.

While flotation is the most commonly used for detection and quantification of coccidia (Hobbs et al., 1999; Hu et al., 2016; Rinaldi et al., 2011), we here used a complementary approach of flotations and diagnostic PCR. We observed relatively large discrepancies between both methods. Flotation and counting of oocysts has a relatively high limit of detection (Ballweber et al., 2014; Webster et al., 1996), explaining negative findings in oocyst flotations positive for PCR. Negative PCR results for samples with visible oocyst in flotations could be a result of a failure to break oocyst walls during DNA extractions and/or faecal PCR inhibition (Raj et al., 2013). Importantly, tested but couldn’t find any species-specific bias in both methods making e.g. relative species prevalences reliable.

Traditional identification of *Eimeria*, depends on the expertise to recognise the morphology of sporulated oocyst (Levine and Ivens, 1965). We show that interpretation of morphometrical data is complex due overlap between species while measurement means agree with literature (Table 1). Considering the challenges of identification and characterisation of *Eimeria* isolates from field samples, we conclude that characterisation of *Eimeria* species requires molecular markers and phylogenetic analysis.

Sequence identity of our isolates to reference sequences from *Eimeria* species previously described in *M. musculus* was above 98% for COI, which is sometimes assumed sufficient correspond differences within species of *Eimeria* (Yang et al., 2015). We confirm taxonomic assignment based on highly supported maximum likelihood and Bayesian phylogenetic clustering of 18S and COI sequences. Moreover, the three identity of the three *Eimeria* species was supported by phenotypic characteristics: morphometry of oocysts and tissue occurrence of the infection.

For some *Eimeria* species precise tissue localization is described based on histology or electron microscopy (Ankrom et al., 1975; Šlapeta et al., 2001). Both methods provide detailed information on developmental stages, but are also time consuming and require a high level of expertise. As an alternative to determine (only) the rough occurrence of the infection along the proximal-distal axis of the intestine, a DNA based qPCR method allowed us not only to detect the presence of *Eimeria*, but also to estimate tissue specific intensity of infection. The qPCR targets a single-copy nuclear gene from the host and a mitochondrial gene of the parasite present in multiple copies (up to 180; Heitlinger et al 2014) to increase sensitivity for *Eimeria* detection.

While infection with rodent *Eimeria* in general can be limited to the duodenum and jejunum (Kvičerová et al., 2007), house mouse *Eimeria* have been described to be mostly found in either the small or the large intestine (Ankrom et al., 1975; Ernst et al., 1971; Haberkorn, 1970; Levine and Ivens, 1965). Using ileum, as the most distal part of the small intestine, and cecum, as the most proximal of the large intestine, we aimed to provide the most stringent test for differences in the site of infection possible: strong infections can be expected to spread in the neighboring tissue, but one could still expect the primary tissue to be more strongly infected. Additionally, genotyping DNA derived from these tissue allowed to detect double infections with *E. vermiformis* in the small intestine and *E. ferrisi* in the large intestine. Localization generally agrees with previous descriptions for the isolates we identified as *E. ferrisi, E. falciformis* and *E. vermiformis* by phylogenetic clustering. Reports of co-infections has been done previously in *A. sylvaticus* from the same colony (Higgs and Nowell, 2000) or in large populations of grey and red squirrels (Hofmannová et al., 2016). To our knowledge we provide the first report of double infections in wild populations of *Mus musculus*.

Double infections can be problematic for identification of species by genotyping. Simultaneous infection of the caecum with *E. ferrisi* and *E. falciformis* would not be recognized with our qPCR method. We shown that amplification of COI with the commonly used primer pair Cocci_COI (Ogedengbe et al., 2011) is differentially efficient for different *Eimeria* isolates. This primer preference can lead to a misidentification in double infections due to the generation of “chimeric” isolates that present different and contradictory information for different markers. Attention to such discrepancies is needed when collating database sequences and when developing multi-marker approaches in general. For rodent *Eimeria* systems, we develop an alternative COI primer pair and find no evidence for differential amplification bias in our cross-validation of the different primer pairs.

*E. ferrisi* is by far the most prevalent species in our study area infecting *M. musculus*. Concerning the house mouse hybrid zone we find infections in *M. m. domesticus, M. m. musculus* and hybrids, suggesting that there are no strict geographical or host subspecies constraints for this species. Population structure for *E. ferrisi* (which could in turn correspond to host subspecies (Goüy de Bellocq et al., 2018; Kváč et al., 2013) can not be found at the resolution the analysed markers provide.

We found that prevalences of *E. ferrisi* and *E. falciformis* increase with increasing host density at the level of farms. This is in agreement with predictions from epidemiology that in large and dense populations contact rates increase (Tompkins et al., 2011) and microoganisms with direct transmission become more prevalent. Such prevalence – host density relationships have been well documented for Hantavirus infections in Bank vole *(My. glareolus)* (Adler et al., 2008; Khalil et al., 2014; Sauvage et al., 2003). In free-living populations of house mice increased host density has been observed to result in higher prevalences of Murine Cytomegalovirus (MCMV) (Singleton et al., 2000). For eukaryotic parasite the prevalence of cestodes and nematodes has been described to be host density dependent in wild and laboratory rodents (Haukisalmi and Henttonen, 1990; Scott and Lewis, 1987). We here document such host density – prevalence relationship for the first time at a species level in *Eimeria* of house mice.

We suggest that species level identification of parasites in wildlife systems will help to assess such questions in more detail and is absolutely required for other questions. For example virulence-prevalence trade-off (Anderson and May, 1982; Frank, 1996) can only be assessed at the species level. In our system one would predict a lower virulence for the prevalent *E. ferrisi* compared to *E. falciformis* and *E. vermiformis*. We have indications from laboratory experiments that such a lower virulence of *E. ferrisi* might be observed compared to *E. falciformis* (Al-khlifeh et al., 2019), while contrary results have been reported before (Tilahun and Stockdale, 1981). We consider this an observation warranting further research.

In conclusion we argue that *Eimeria* in wildlife populations should be identified more frequently at the level of species previously described by taxonomists. We propose to integrate a set of simple methods into a reproducible procedure to achieve this aim. For Coccidians, as important parasites of vertebrates, only species specific assessment will allow to test hypotheses in evolution, ecology and epidemiology.

## Acknowledgements

The authors acknowledge our collaboration partners Prof. Dr. Jaroslav Piálek and his team from the Institute of Vertebrate Biology, AS CR, Brno, Department of population Biology in Studenec, Czech republic for their valuable help during the collection of samples. To Deborah Dymke, Thi Phuong Le and Julia Murata for their assistance during the processing within Heitlinger Group.

