## Supplementary figures and images for "Detection and quantification of house mouse *Eimeria* at the species level – challenges and solutions for the assessment of Coccidia in wildlife"

### Supplementary data S3

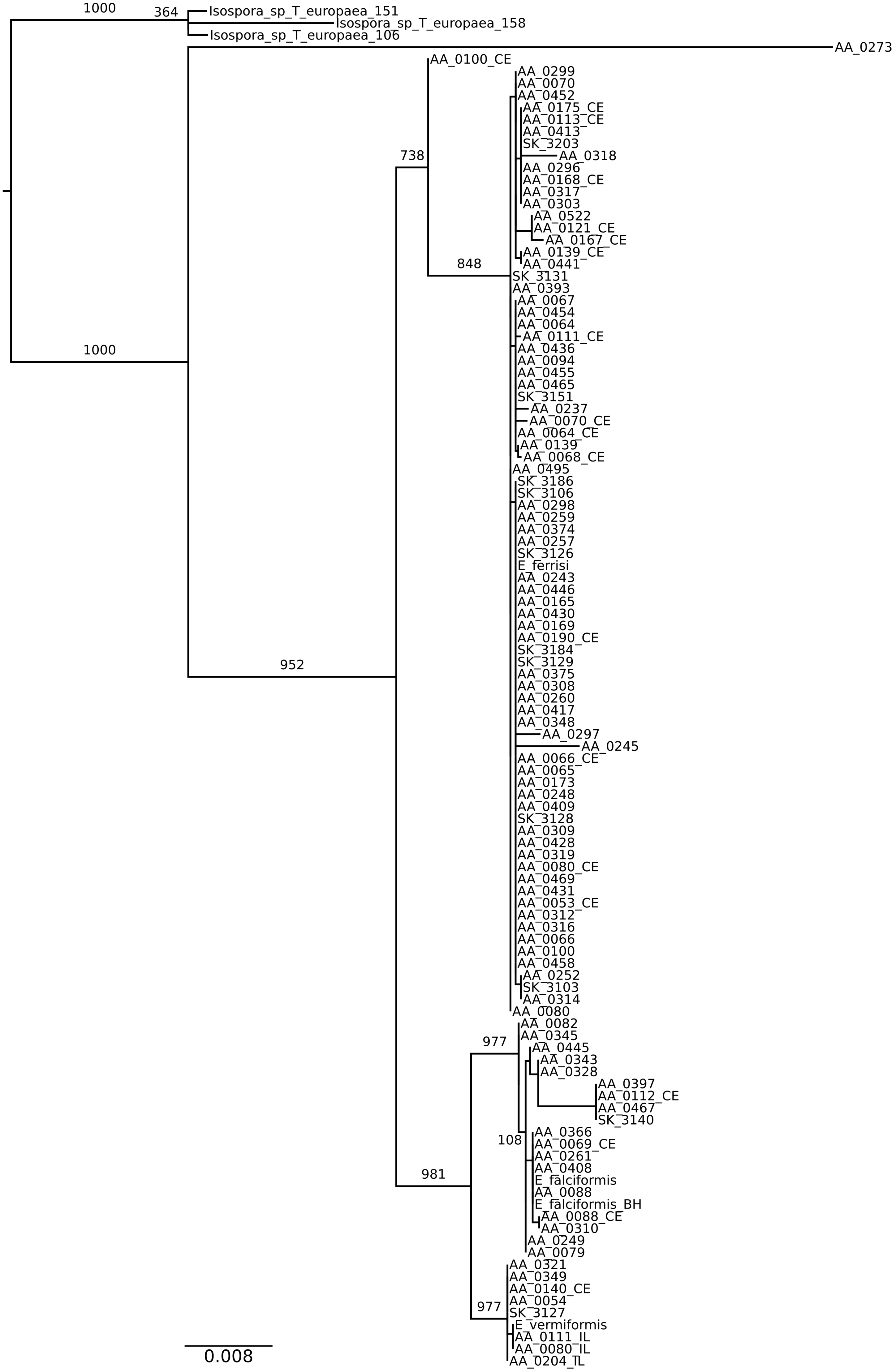

### Supplementary data S4

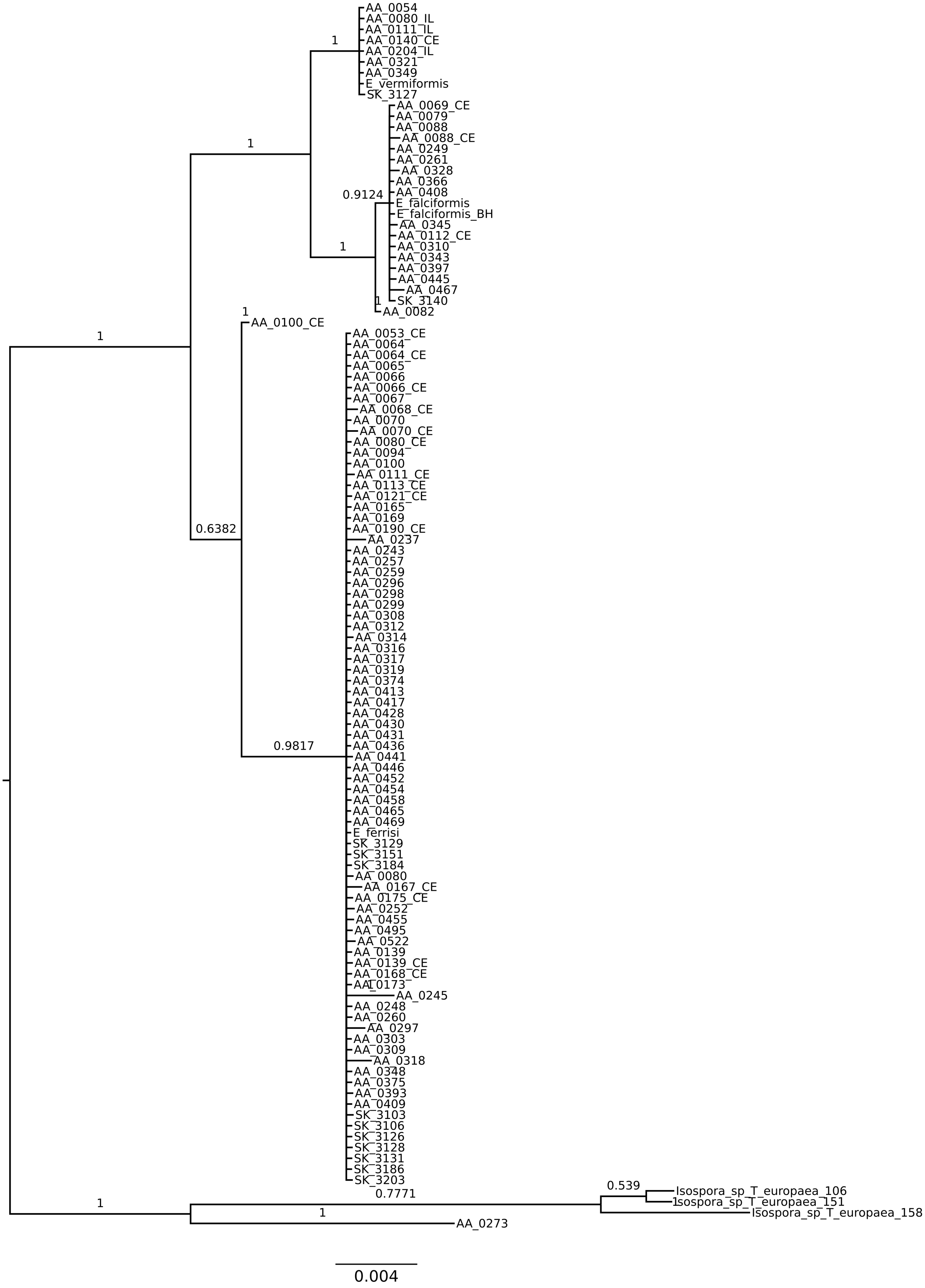
